# Preparing glycomics data for robust statistical analysis with GlyCompareCT

**DOI:** 10.1101/2022.05.31.494178

**Authors:** Yujie Zhang, Sridevi Krishnan, Bokan Bao, Austin W.T. Chiang, James T. Sorrentino, Song-Min Schinn, Benjamin P. Kellman, Nathan E. Lewis

**Affiliations:** Department of Pediatrics, University of California, 9500 Gilman Drive MC 0760, La Jolla, San Diego, CA 92093, USA; Department of Biostatistics, Harvard T.H. Chan School of Public Health, 677 Huntington Avenue, Boston, MA 02115, USA; Department of Bioengineering, University of California, 9500 Gilman Drive MC 0760, La Jolla, San Diego, CA 92093, USA

## Abstract

**Summary:** Glycomics data are rapidly increasing in scale and diversity. Complexities in glycan biosynthesis (hierarchy, competition, and compartmentalization) make preprocessing critical to address resulting sparsity (many similar glycosylation profiles may share few common glycans) and non-independence (substrate-competition in glycan biosynthesis results in non-independence incompatible with many statistical methods). Here, we present GlyCompareCT, a portable command-line tool, to address these challenges thereby facilitating downstream analyses. Given glycan abundances, GlyCompareCT conducts substructure decomposition to quantify hidden biosynthetic intermediate abundance and relationships between measured glycans. Thus, GlyComparCT mitigates sparsity and makes interdependence explicit thereby increasing statistical power. Ultimately, GlyComparCT is a user-friendly implementation of substructure analysis designed to increase accessibility, interoperability, and scope and consistency in glycomics analysis.

**Availability and implementation:** Source code, test data, and instructions for GlyCompareCT v1.1.0 are available at: https://github.com/LewisLabUCSD/GlyCompareCT

**Supplementary information:** https://github.com/LewisLabUCSD/GlyCompareCT/raw/main/Supplementary%20Material.pdf

## Introduction

Glycosylation, the addition of branched sugar polymers called glycans, occurs on a variety of biomolecules. Glycans are composed of multiple monosaccharides linked by glycosidic bonds, which enable tremendous diversity of structure through differences in linkage stereotype and carbon positions. However, many glycans contain common substructures, which reuse the same monosaccharides and linkages. Glycosylation is a hierarchical process complicated by numerous processive, and high-specificity biosynthetic enzymes that compete for common substrates^1^. Innovations in mass spectrometry enabled rapid growth in the number of discovered glycans.^2–11^. Comprehensive, and consistent analytics are increasingly critical as rich glycoprofile datasets are cataloged into ever-larger datasets and databases^12–18^. Sparsity of glycan profiles across observations and structural commonalities between different glycans (non-independence) create challenges in rapid and accurate comparison of glycoprofiles. GlyCompare^19^ is a Python library, effective across multiple glycan-types^20–26^, to decrease sparsity and specify non-independence in glycomics data, thereby enabling large-scale analytics and comparison.

Shared glycan substructures, arising from common biosynthetic intermediaries, are often neglected in whole-glycan analysis. Consequently, glycans differing by only a single monosaccharide can be considered as independent and distinct structures. Considering these glycans to be distinct ignores their biosynthetic history and corresponding inter-dependencies. Both singular and cascading biosynthetic shifts, overlooked by traditional glycan analysis, become evident when comparing substructure abundance profiles.

To improve accessibility and consistency for the central GlyCompare functionality (substructure decomposition), we introduce GlyCompareCT, a command-line wrapper of GlyCompare intended for convenient, portable, and accessible command-line usage.

## Interpretation

GlyCompare calculates the latent abundance of biosynthetic intermediates by decomposing glycans into substructures. Glycan abundance profiles, glycoprofiles, are typically created through software-aided^27–29^ assembly from spectrometry-fragmented glycan compositions or retention-time matching in chromatography. While spectrometry data is already fragmented, the fragmentation biases of each pipeline are not biosynthetically relevant. Therefore, substructure abundances are most interpretable and interoperable when calculated from completed glycoprofiles. A substructure abundance is calculated either as (1) the sum of abundances over all glycans containing that substructure (binary multiplier), or (2) the glycan abundance sum after scaling each glycan abundance by the number of substructure occurrences in that glycan (integer multiplier). Substructure abundance is therefore a proxy for precursor or upstream abundance; the total amount of a substructure ever made in that profile. Glycomotifs are the minimal set of the largest non-redundant substructures; sufficient to describe all variance in the dataset. The user can specify a “biosynthetic” reducing-end substructure (e.g., N-, O-, lactose, or custom) or an “epitope” non-reducing-end substricture (e.g., all monosaccharides, or Sialyl Lewis A) root from which to search for the shared substructures. Finally, representative substructures can be calculated to summarize substructure clusters in the substructure-abundance bicluster. GlyCompare-enabled substructure analysis simplifies glycome analytics and increases statistical power by decreasing sparsity and specifying non-independence. (Figure 1a)

**Figure 1.**
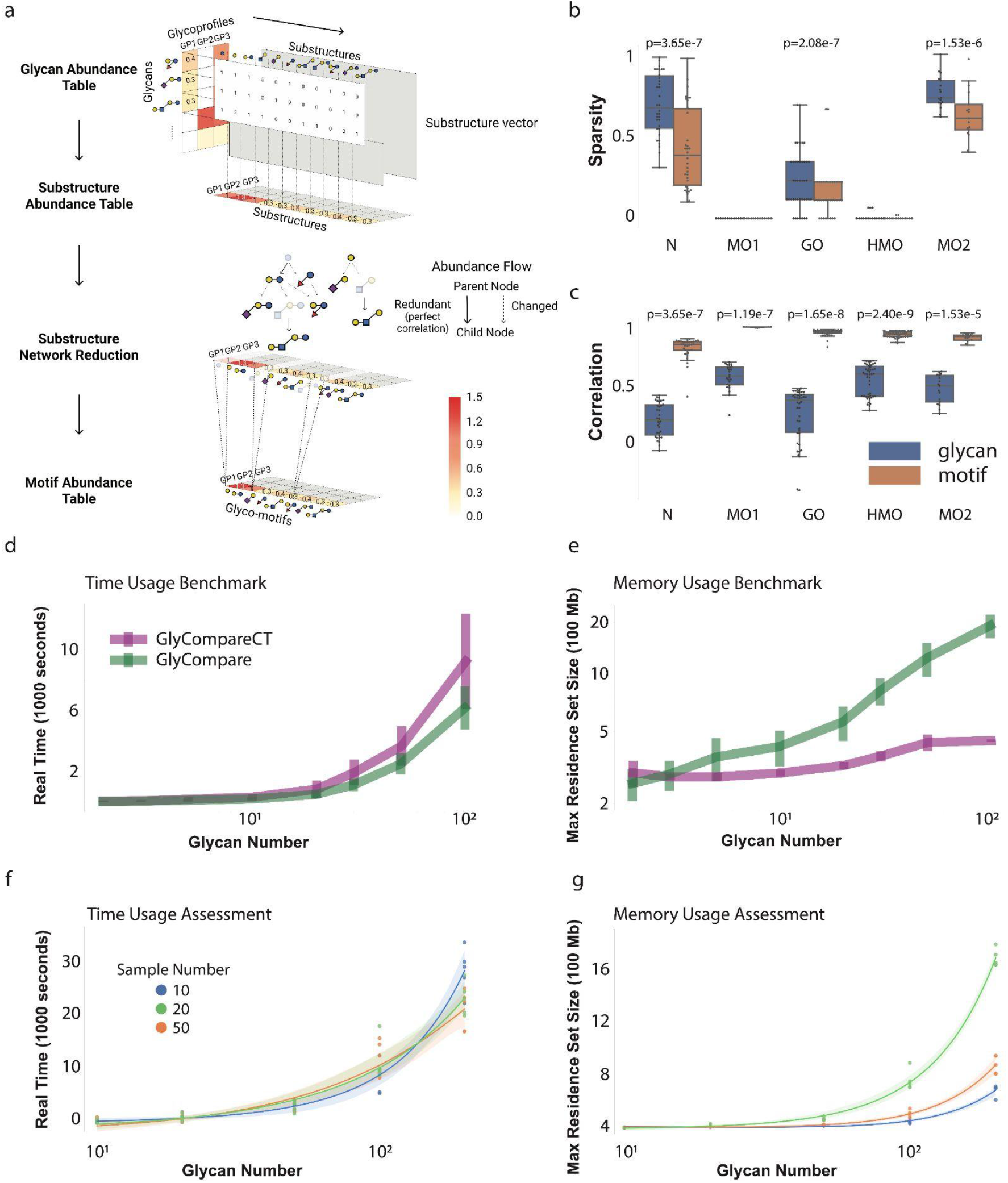
GlyCompareCT decomposes measured glycans into their substructures and extracts motifs, to decrease sparsity and capture interdependence of measured glycans. (a) GlyCompareCT decomposes glycans to their substructures and generates a substructure vector wherein each entry is the frequency of substructure within the corresponding glycan. GlyCompareCT subsequently calculates the substructure abundance table through multiplication of the glycan abundance table with the substructure vector. A substructure network is constructed to reduce perfectly correlated substructures. Only leaf substructures are retained among perfectly correlated substructures. Finally, GlyCompareCT outputs the remaining entries as a glyco-motif abundance table. (b, c) GlyCompareCT was tested on erythropoietin N-glycosylation (N^21,22^), human milk oligosaccharides (HMO^20^), mucin-type O-glycans (MO1^24^, MO2^23^), and gangliosides (GO^25^) with default parameters (see **Implementation**). (b) Boxplots show that glycoprofile sparsity (zero-abundance fraction) decreased in three of five datasets after substructure decomposition to glycomotif-profiles. (c) Boxplots show inter-profile correlation increases between glycomotif profiles after substructure decomposition. Boxplots (b-c) display the median (center-line), interquartile range (IQR; box), and 1.5*IQR (whiskers). (d, e) Explore performance improvements in GlyCompareCT v1.0.0 over Glycompare v1.1.3 (see **Performance**). Line plots with error bars (standard deviation) indicate minimal changes to runtime (d) and substantial improvements in memory (e). (f, g) Time and memory usage performance on varying numbers of glycans and samples. Each pair of (glycan_num, sample_num) contains five trials. Line plots are linear regression lines fitted to runtime (f) and memory (g).

## Performance

Full GlyCompareCT methodology description can be accessed at https://github.com/LewisLabUCSD/GlyCompareCT/blob/main/Methodology%20Supplementary.pdf

GlyCompareCT, like GlyCompare, is designed to mitigate sparsity (zero-abundance fraction) while identifying non-independent glycan substructures across diverse datasets. We tested GlyCompareCT on N-link, O-link, structural and compositional datasets (Figure 1b-c) with default parameters. In three of five datasets, glycomotif abundance tables were less sparse than corresponding glycan abundance tables. In the remaining two datasets, glycan abundance sparsity was already close to zero (an artifact of targeted measurement of select glycans). Additionally, correlation between glycomotif profiles was higher than correlation between glycan abundance profiles across all datasets. We tested all combinations of all parameters and found that regardless of parameterization, sparsity decreased (Wilcoxon test, p ≤ 6.70e-4) and correlation increased (Wilcoxon test p ≤ 1.53e-5).

Notably, modifications to GlyCompare v1.1.3 (substructure-occurrence matrix representation) resulted in substantial improvements to memory performance in GlyCompareCT v1.0.0; particularly with large glycomic datasets. We compared GlyCompare and GlycompareCT using unit-datasets (all glycan abundances are 1, each dataset contains 1 sample, and 2, 3, 5, 10, 20, 30, 50, or 100 glycans chosen randomly from the linkage-specific glycomotif reference [github.com/yuz682/GlyCompareCT/tree/main/reference]). We analyzed 80 unit-datasets, 10 datasets for each dataset size. GlyCompareCT showed a marginal but significant time increase compared to GlyCompare (1.5-1.7 fold; 2-sample 2-sided t-test p-value<0.016, Figure 1d). The maximum resident set size (RSS) benchmark between GlyCompareCT and GlyCompare showed memory usage substantially decreased with GlyCompareCT. Analyzing unit-datasets with 100 randomly selected glycans, GlyCompareCT approaches a 5-fold reduction in memory (from 1923Mb to 446Mb, Figure 1e). With a small increase in runtime, GlyCompareCT decreased memory usage from exponential to linear as a function of glycan count.

To further assess the runtime and memory performance of GlyCompareCT, we tested it on a series of paired data varying glycan number and sample number. Glycan numbers are 10, 20, 50, 100, and 200. Sample numbers are 10, 20, and 50. For each pair of glycan number and sample number, we ran five trials. For each trial, we chose glycans randomly and randomly set 80% of entries to 0 and the rest to 1 to simulate the sparsity in real glycoprofiles. The time usage does not vary significantly across different sample numbers (Figure 1f) while the memory usage increases with more samples (Figure 1g).

## Conclusion

GlyCompareCT allows fast and accurate glycoprofile comparisons and increases the statistical power for glycomic data analysis. As a command line tool extension of the GlyCompare python package, GlyCompareCT improves usability and consistency necessary to aid analysis of growing glycomics datasets.

## Supporting information

Methodology Supplementary

## Acknowledgements

This work was conducted with generous support from NIGMS (R35 GM119850), NHLBI (K12HL141956) and the Novo Nordisk Foundation through the National Biologics Facility at the Technical University of Denmark (NNF20SA0066621).

