## Supplementary material for "Preparing glycomics data for robust statistical analysis with GlyCompareCT": Methodology Supplementary

**Methods**

GlyCompare decomposes glycan structures into substructures and then identifies all glycomotifs necessary to describe the glycoprofiles provided. GlyCompare additionally generates a bi-clustered heatmap displaying glycomotif relative abundance across glycoprofiles, and glycomotif clusters are summarized by representative substructures (abundance aggregation of glycomotifs containing a representative substructure). (Figure 1a)

**Substructure Identification and Decomposition**

For each glycan, GlyCompare breaks one or multiple glycosidic bonds to permute all possible substructures. GlyCompare then sorts the substructures’ monosaccharide counts. Each substructure is annotated for occurrences in each glycan. Glycosidic linkage information can be leveraged (“linkage-specific”) or ignored (“structural”) during substructure extraction depending on the application. Substructures are named by their index position in the glycomotif vector (hosted at https://github.com/LewisLabUCSD/GlyCompareCT/reference); novel glycomotifs will be appended to the end of the glycomotif vector.

For each glycan in a glycoprofile, all substructures are enumerated (substructure-occurrence matrix). Next, GlyCompare constructs a substructure abundance table from the glycoprofile. Each substructure can appear in multiple glycans. The abundance of each substructure within the glycoprofile is calculated by summing over the products of each glycan abundance and the substructure occurrences in that glycan. Suppose the substructure occurrence of substructure *i* in glycan *j* is denoted as *o_ij_* and the abundance of glycan *j* is *a_j_* , then the abundance of substructure *i* in the glycoprofile denoted as *s_i_* is *s_i_*=*Σ_j_o_ij_a_j_* . The substructure abundance values for each substructure across glycoprofiles represent the “decomposition” of glycoprofiles into substructure abundance profiles.

**Glycomotif extraction from the substructure network**

To identify glycomotifs, GlyCompare identifies related substructures and quantifies substructure abundance correlation. Related substructures are retained only if they are not correlated. Specifically, when traversing the substructure network from a user-specified root(s), the smaller substructure of two perfectly correlated substructures is dropped.

Glycan substructures relations are represented in a directed acyclic substructure network where nodes are glycan substructures and edges directionally connect substructure nodes separated by the addition of one monosaccharide; therefore, edges represent known or hypothetical glycosynthesis reactions. Each node can have multiple parent nodes and multiple child nodes as many glycans are substrate to multiple biosynthetic reactions each yielding distinct glycan products. Users specify the substructure network root as either each monosaccharide (epitope mode) or a specific biosynthetic root; specifying the first structure examined helps focus the network reduction on biosynthetic or epitope motifs of interest.

Next, the nodes and edges in the substructure are annotated with substructure abundance and correlation respectively. These annotations guide the removal of redundant substructures. Each substructure node is annotated with the corresponding substructure abundance vector–abundance of that substructure across samples. Each edge is annotated with the correlation between the substructure abundance vectors connected by that edge.

The correlations now associated with each edge guide the removal of redundant substructures. Redundant substructures, those sharing an edge in the substructure network and which are perfectly correlated across samples, are reduced to the largest (child) substructure thereby compressing the network without information loss. We call the minimal substructure set ‘glycomotifs’, and the corresponding subset of substructure abundances is the glycomotif abundance table.

**Glycomotif profile clustering**

GlyCompareCT will optionally visualize the glycomotif abundance table as a hierarchically bi-clustered heatmap; clustering similar abundance vectors (profiles) and glycomotifs together. Groups of similar glycomotifs can, optionally, be summarized as a summary or “representative” substructure–a glycan substructure containing all monosaccharides included in more than half of in-group glycomotifs.
